# Missense variants reveal functional insights into the human ARID family of gene regulators

**DOI:** 10.1101/2021.11.17.468850

**Authors:** Gauri Deák, Atlanta G. Cook

## Abstract

Missense variants are alterations to protein coding sequences that result in amino acid substitutions. They can be deleterious if the amino acid is required for maintaining structure or/and function, but are likely to be tolerated at other sites. Consequently, missense variation within a healthy population can mirror the effects of negative selection on protein structure and function, such that functional sites on proteins are often depleted of missense variants. Advances in high-throughput sequencing have dramatically increased the sample size of available human variation data, allowing for population-wide analysis of selective pressures. In this study, we developed a convenient set of tools, called 1D-to-3D, for visualizing the positions of missense variants on protein sequences and structures. We used these tools to characterize human homologues of the ARID family of gene regulators. ARID family members are implicated in multiple cancer types, developmental disorders, and immunological diseases but current understanding of their mechanistic roles is incomplete. Combined with phylogenetic and structural analyses, our approach allowed us to characterise sites important for protein-protein interactions, histone modification recognition, and DNA binding by the ARID proteins. We find that comparing missense depletion patterns among paralogs can reveal sub-functionalization at the level of domains. We propose that visualizing missense variants and their depletion on structures can serve as a valuable tool for complementing evolutionary and experimental findings.

## Introduction

Advances in high-throughput sequencing have led to a sweeping expansion in genetic variation data of human protein-coding genes. With nearly 15 million curated exome variants made available by the international Genome Aggregation Database (gnomAD)(1), statistical analyses have identified genes that are intolerant to loss-of-function (LOF) and are likely associated with disease (1, 2). This has aided large-scale assessments of genetic causality in autism spectrum disorder (3) and inherited cardiomyopathies (4), and has also led to novel frameworks for evaluating drug toxicity or efficacy through consideration of the genetic constraint on target genes (5, 6). The increase in statistical power afforded by the size of these datasets has allowed genes to be ranked for their importance to human health based on their intolerance to variation.

Beyond the effects of LOF variants on a gene, constraint analyses can be further narrowed to focus on missense variants at the level of a protein domain. We define a domain as an independent folding unit or a conserved sequence block that is likely to approximate an independent folding unit. Depletion of missense variants in whole domains, or specific segments within domains, has been found to correlate with evolutionary conservation (7). Rare mutations occurring in such regions are likely to be pathogenic (8, 9). In agreement, saturation mutagenesis studies have shown that mutation-intolerant regions map to conserved protein domains, particularly at residues that are involved in DNA, protein, or ligand binding and are associated with pathogenic variants (10, 11). In another study, domain-level definitions improved disease prediction scores compared to random segmentation of sequences (12). These findings indicate that, like whole genes, functionally important regions in proteins are subject to negative selection. Patterns in population-wide missense variation could therefore be harnessed to gain insights into protein function (13).

While calculating variant distributions along protein sequences can help to identify essential domains, it fails to consider the arrangement of variants in 3D space (8). This has been addressed by manual mapping of variants (14) or computational mapping of variant depletion scores directly onto protein structures. For example, Hicks et al., developed a ‘3D Tolerance Score’ which compares an observed and expected number of missense variants in 5 Å-radius spheres around individual atoms in a 3D structure (15). In an alternative approach, Tang et al. introduced PSCAN, which scores the spatial dispersion of missense variants onto structures (16), based on a previous finding that neutral variants tend to be dispersed, while pathogenic variants cluster (17). In a further approach, the MISCAST suite provides an approach to connect probabilities of loss of function with primary sequence level features (18).

Collectively, the above studies show that surfaces of proteins where important functional sites are located are depleted of missense variants. Statistical approaches enable sorting and/or predicting functional sites using proteome-wide approaches but may not necessarily enable non-specialists to inspect an individual protein of their choice. We developed two programs to allow easy visual inspection of variants on primary sequences (1D) and tertiary structures (3D) from the gnomAD database (Fig. 1A).

**Figure 1:**
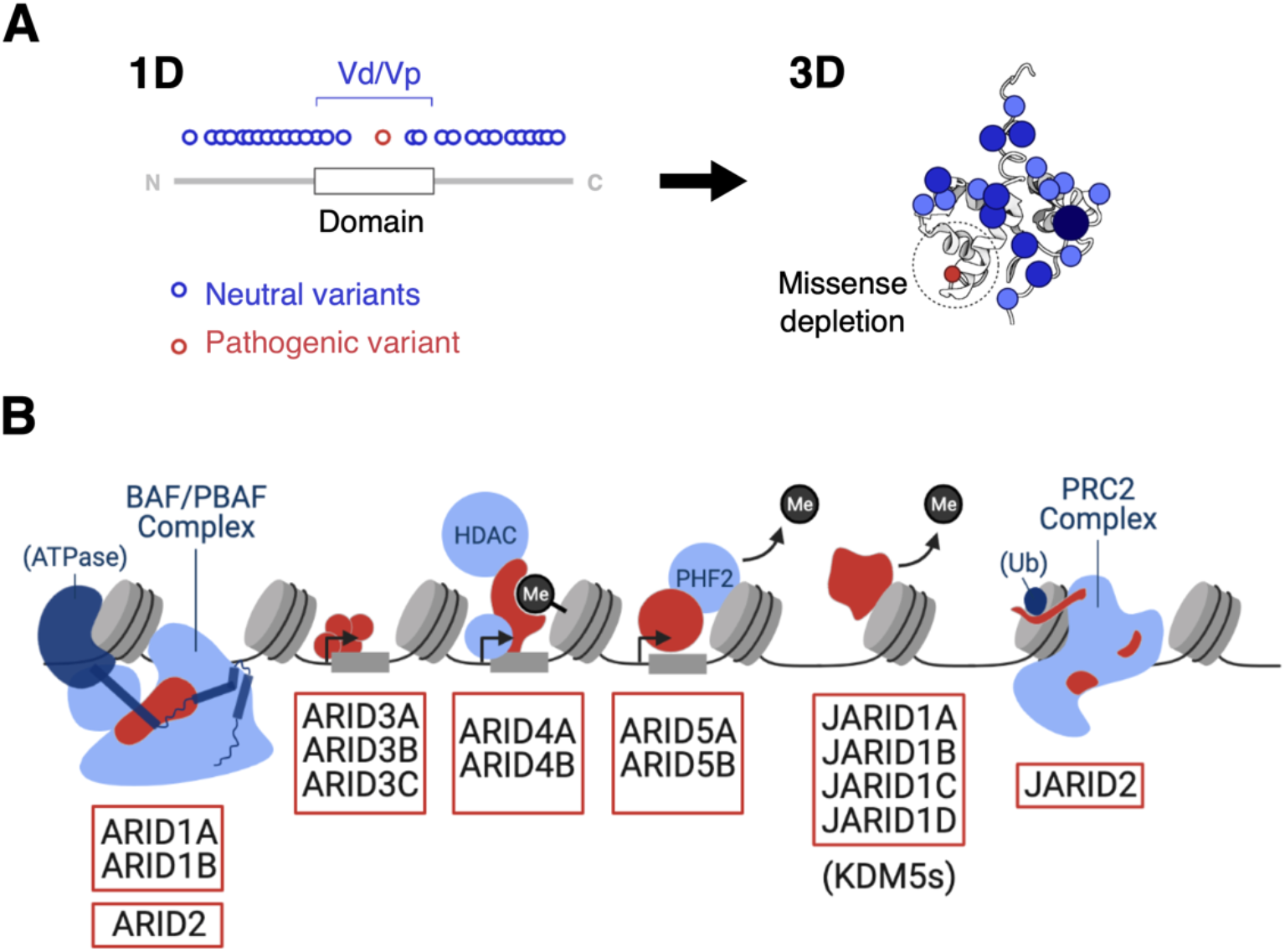
**(A)** Schematic of the 1D-to-3D approach of mapping missense variants onto protein structures. **(B)** Schematic illustration of the interactions of ARID family members (red) with nucleosomes (grey) and other chromatin-binding proteins (light blue).

First, we generated a program to calculate the average density of missense variants in a protein domain (Vd) to the average density of missense variants in the whole protein (Vp). We show that plotting variants in 1D together with the ratio of Vd/Vp helps identify domains that are missense depleted. The standalone values of Vp for the ARID family members exhibit good correlation with the missense Z score (mZ), typically used in gnomAD (2).

Second, we developed a simple, convenient program called 1D-to-3D to map the same variant data onto any 3D structure, representing variants as spheres of increasing size and color intensity with increasing allele frequency (AF) (Fig. 1A). This allows users to find surfaces of the protein that are depleted for missense variants and to compare this with complementary information such as surface conservation.

Using 1D-to-3D, we performed a comprehensive analysis of missense variation in the “<u>A</u>T-<u>r</u>ich <u>i</u>nteractive <u>d</u>omain” (ARID) family of gene regulators (Fig. 1B). This family was selected due to the clinical significance of its members in multiple cancer types (19–24), rare developmental disorders such as Coffin-Siris syndrome (25), and immunological diseases including systemic lupus erythematosus, type II diabetes, and atherosclerosis (26–28). In humans, the ARID family comprises 15 proteins that can further be categorized into 7 subfamilies, all of which have an ARID DNA binding domain (29). Despite their common domain, the family members regulate distinct sets of genes via diverse molecular mechanisms (Fig. 1B). ARID1 and ARID2 are core subunits of the BRG1/BRM-associated factors (BAF) family of nucleosome remodeling complexes while JARID2 is an accessory subunit of the Polycomb Repressive Complex 2 (PRC2) (30–34). In contrast, ARID3 proteins are transcription factors (35), while ARID4s and ARID5s are adapter proteins that recruit other transcriptional regulators such as the mSin3A-histone deacetylase (HDAC) complex and PHD Finger 2 (PHF2) histone demethylase respectively (36–38). ARID5A is thought to be an RNA-binding protein (39, 40). Finally, the four JARID1 proteins are enzymes that mediate transcriptional changes by removal of histone H3K4 di-/tri-methylation marks (21).

Here we provide a comprehensive analysis of the ARID family as a whole, including domain architecture mapping, phylogenetic analysis and searching for known pathogenic variants. We complemented these analyses with our 1D-to-3D approach to identify surfaces of proteins that are depleted (or not) of missense variants, to provide a deeper annotation of functional sites within these proteins.

## Materials and Methods

### Sequence Alignments and Domain Annotations

Sequences of ARID family orthologs were selected using the Oma orthology database (RRID:SCR_016425) (41). Multiple sequence alignments were generated using MAFFT (RRID:SCR_011811) (42) and pairwise alignments were generated with EMBOSS Needle (43). Alignments were then visualized in JalView 2.11.1.4 (RRID:SCR_006459) (44). Evolutionary relationships between paralogs in ARID subfamilies were verified using TreeFam (RRID:SCR_013401) (45). A structural sequence alignment for the ARID5B BAH domain was created using the DALI server (RRID:SCR_013433) (46). Functional domains of each family member were annotated using InterPro (47) or based on experimental data (the ARID1A/1B core binding regions (48), ARID4A/B R2 region (37), and JARID1A (49) and JARID1B (50) domains).

### Structures and Models

A complete list of analyzed structures is in Supplementary Table 1. For domains with no available structure, we used models from the AlphaFold protein structure database (51). Only regions with predicted local-distance difference test (pLDDT) scores > 70 were considered. The pLDDT score is a confidence measure that reflects the validity of local inter-atomic distances in a predicted structure. A cut-off of > 70 is considered a “generally correct backbone prediction” (52). All of the structures were visualized in PyMOL (53). Surface electrostatics scores for the ARID4A and ARID4B hybrid Tudor domains were calculated using the Adaptive Poisson-Boltzmann Solver (RRID:SCR_008387) (54) in PyMol.

### Surface Conservation

Surface conservation was mapped onto the structures of ARID1A (PDB ID 6LTJ, chain L), ARID1B (AlphaFold model, UniProt ID: Q8NFD5, amino acids (aa) 1593-1699 and 1905-2236), ARID2 (AlphaFold model, UniProt ID: Q68CP9, aa 155-464), JARID1A (PDB ID: 5CEH), and JARID1B (PDB ID: 5FUP) using ConSurf (RRID:SCR_002320)(55). For ARID1A/1B, MAFFT alignments of 70 Oma group vertebrate orthologs were submitted to the server. For ARID2 and JARID1A/1B, MAFFT alignments of 165 and 80 Oma group metazoan orthologs were submitted respectively. We selected Oma groups because they exclude paralogs and include only one co-ortholog if several are found for a given species. This yields a collection of non-redundant sequences that can be filtered at specific taxonomic levels (56). All sequence alignments can be accessed at: https://doi.org/10.7488/ds/3190.

### Isoform expression analysis

Isoform-specific expression data for ARID5B was obtained from ISOexpresso (57). The ratio of Isoform 1 (uc001jlt.2) to Isoform 2 (uc001jlu.2) expression levels was compared across 735 samples from 22 human tissue types of healthy individuals.

### Constraint Metrics

Established constraint metrics including pLi, LOEUF, missense Z, and RVIS scores for each ARID family member were obtained from the official gnomAD and Genic Intolerance web browsers (1, 58). P-values corresponding to missense Z-scores were calculated using the Excel NORM.S.DIST function (the output for positive Z-scores was subtracted from 1). A complete list of collected metrics is available in Supplementary Table 1.

### Variant Data Processing

7,652 non-synonymous variants associated with UniProt canonical sequences of the 15 ARID family proteins were extracted from the gnomAD v2.1.1 dataset (GRCh37/hg19) (1). The dataset is publicly available and contains variants from 125,748 quality-controlled exomes of unrelated, adult individuals not affected by severe pediatric disease (1). To ensure our analysis was restricted to neutral missense variants, only variants with the Variant Effect Predictor annotation ‘missense’ were considered, and variants with the ClinVar annotation ‘pathogenic’, ‘likely pathogenic’, ‘conflicting interpretations of pathogenicity’, and ‘uncertain significance’ were filtered out. To perform this filtering, we developed a Python program called 1D-to-3D.py, which processes variants from csv files downloaded directly from gnomAD (1) (further details in “3D Visualization”). After filtering, 7,540 variants were used for further analyses. All raw and processed gnomAD data can be accessed in supporting information Supplementary Table 2 and Supplementary Table 3 respectively.

### Pathogenic Variants

A family-wide search for pathogenic missense variants was performed using ClinVar (RRID:SCR_006169) and DECIPHER (RRID:SCR_006552), two publicly-accessible databases of clinical variants and their phenotypes (59, 60). Using ClinVar, we identified variants with the search criteria ‘missense’ and ‘pathogenic’ or ‘likely pathogenic.’ In DECIPHER, we searched for ‘research variants’ from the Deciphering Developmental Disorders Study, which collected variants from ~14,000 UK children with undiagnosed developmental disorders (59). All variant accession codes are available in Supplementary Table 1.

### 1D Plots and the Vd/Vp Ratio

Filtered missense variants were mapped onto protein sequences of ARID family members using Plot Protein (61). The Plot Protein R script was modified to allow for color manipulation and domain diagram alterations of the output graphs in Inkscape (62). Vd/Vp ratios of functional domains were calculated using the following formula:

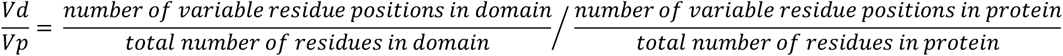

To automate the process, we developed a Python program called VdVp_Calculator.py. Like 1D-to-3D, the Vd/Vp Calculator processes csv files downloaded directly from gnomAD (1). It requires a user-defined text file with domain boundaries and calculates the Vd/Vp ratios of all functional domains in the protein of interest. The program script and user instructions are accessible at GitLab (https://git.ecdf.ed.ac.uk/cooklab/deak).

### 3D Visualization

To visualize missense variation in 3D, the 1D-to-3D program uses filtered data from gnomAD (1) and generates a PyMOL script that maps the variants onto protein structures. The variants appear as spheres at the C*α* of the associated residue and increase in size and shade of blue with increasing AF. In the case of multiallelic sites, the program applies the addition rule for disjoint events, i.e. if multiple variants occur at the same residue position, their AFs are summed. The AF values for each position are compressed using a base 10 log scale and the positions are sorted into 6 bins (AF <10^−6^-10^−5^, 10^−5^-10^−4^…10^−1^-10^0^). Variants in each bin are visualized as spheres of different size and shade of blue. Further details regarding user inputs and numerical handling can be found in the user instructions and program script, accessible at GitLab (https://git.ecdf.ed.ac.uk/cooklab/deak). We used the 1D-to-3D program to annotate 11 solved and 6 modelled structures of the ARID family members with missense variants. A list of these structures, PyMOL selection names, and start/end residues can be found in Supplementary Table 1. The PyMOL scripts can be accessed at: https://doi.org/10.7488/ds/3190.

## Results and Discussion

### Genetic Constraint in the ARID Family

As a preliminary assessment of variation in the ARID family, we collated existing constraint metrics for each member from gnomAD. These included “loss-of-function observed/expected upper bound fraction” (LOEUF) (1) and missense Z (mZ) scores (Fig. 2A) as well as pLi and RVIS scores (Supplementary Table 1) (1, 58). The lower the LOEUF, the fewer the variants observed than expected, indicating negative selection against LOF variation. All ARID family genes except ARID3B/3C and JARID1B/1D can be classified as LOF-intolerant (Fig. 2A). This indicates that they are subject to strong purifying selection and are statistically likely to have disease associations, a higher number of protein-protein interaction partners and broad tissue expression (1). Intolerance to variants in the ARID family is also supported by pLI and RVIS scores (Supplementary Table 1).

**Figure 2:**
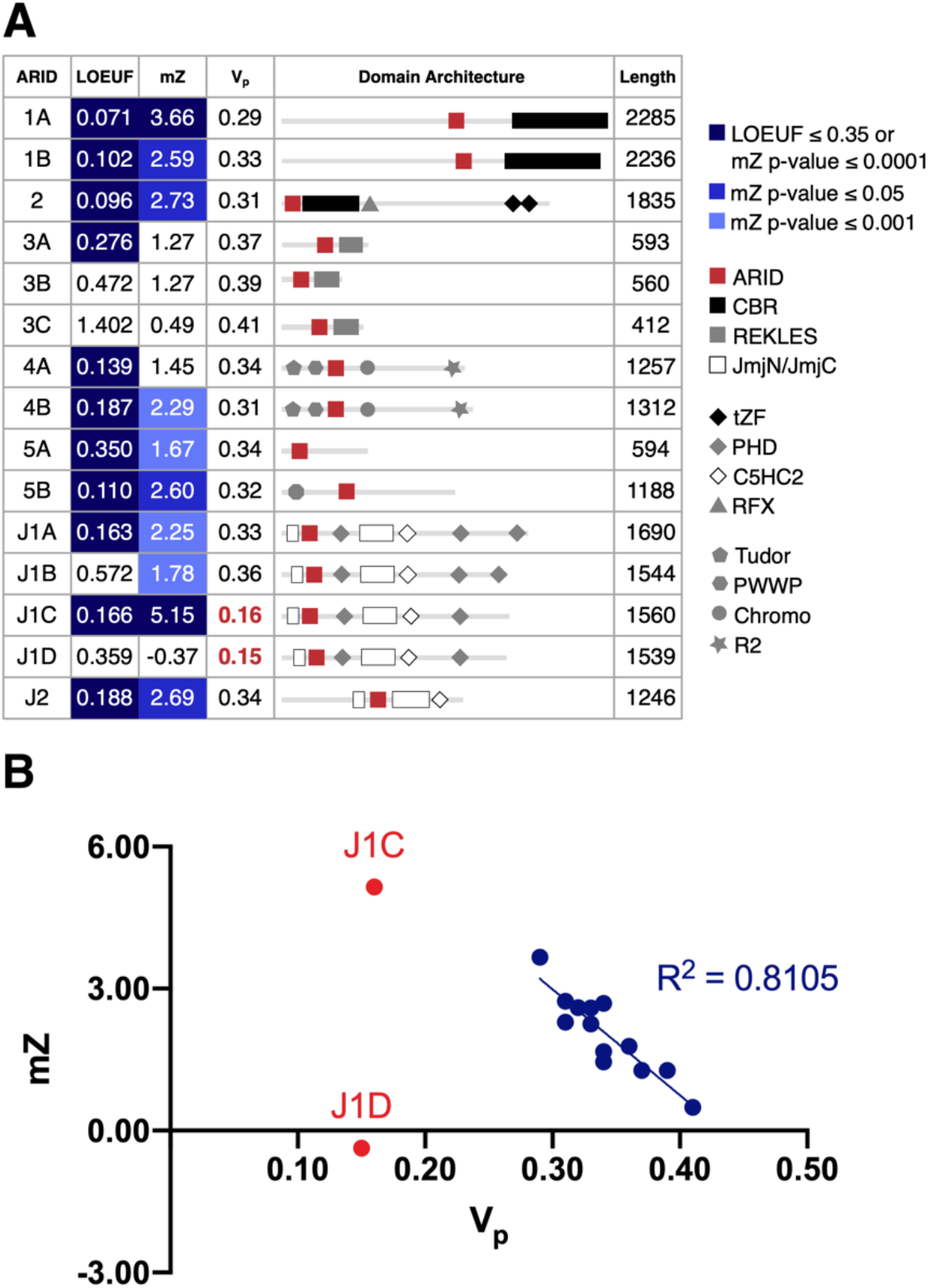
Genetic constraint in the ARID family. **(A)** An overview of constraint metrics, Vps, and domain architecture in the ARID family. LOEUF is the upper boundary of the 90% confidence interval of the observed/expected ratio of loss-of-function (LOF) variants in a given gene. The recommended threshold to segregate LOF-intolerant and LOF-tolerant genes is 0.35. The mZ scores outside of ±3.09 correspond to a recommended P-value threshold of 0.001. **(B)** Correlation between the mZ scores and Vps for this dateset (blue). Values for JARID1C (J1C) and JARID1D (J1D) are red.

The mZ score represents the deviation of the observed from the expected number of missense variants (single amino acid substitutions), for a given gene, where positive scores indicate missense depletion and negative scores indicate missense enrichment (2). ARID1A and JARID1C have mZ P-values of < 0.001 and nearly all members have positive scores, indicating that specific residue positions are also under selective constraint (Fig. 2A).

We calculated Vp values that denote the average density of missense variants for each protein in our dataset. As expected, lower Vp values correlate with higher mZ scores (Fig. 2B). However, JARID1C and JARID1D do not fit the observed correlation between mZ and V_p_ values (Fig. 2B). Rather than missense depletion, this is likely to arise from low data availability because JARID1C and JARID1D are encoded on the X and Y chromosomes respectively (1, 63). This analysis suggests that Vp is a good proxy for mZ and that values outside of the range of 0.29-0.42 may indicate when insufficient data are available for analysis. We excluded JARID1C and JARID1D from subsequent analyses.

These constraint metrics demonstrate that individual ARID family members are under significant selective pressure, yet they are less informative on missense depletion at the level of domains or smaller functional regions (Fig. 2A). To bridge this gap, we developed the ‘1D-to-3D’ approach (Fig. 1A). In ‘1D’, primary protein sequences are annotated with neutral or pathogenic missense variants and Vd/Vp ratios are calculated to compare the linear distribution of missense variants in functional domains. In ‘3D’, the missense variants are mapped onto solved or modelled protein structures. This allows integration of the AFs of variants at each residue position, with visual discrimination of AFs using increasing sphere size and shade of blue (Fig. 1A). In line with larger-scale analyses (7, 15), we hypothesized that the annotated structures would reveal functionally important 3D sites depleted of neutral variants and enriched in pathogenic variants.

### Validating the 1D-to-3D Approach on Known ARID Complex Structures

To verify that the 1D-to-3D approach can be used to identify sites depleted of missense variants for this family, we analyzed the structurally well-characterized ARID1 and JARID1 subfamilies. The ARID1 subfamily comprises ARID1A and ARID1B, two vertebrate paralogs with identical domain architecture (Fig. 2A) and 57 % sequence identity in humans. ARID1A and ARID1B are mutually-exclusive core subunits of the BAF chromatin remodeling complex (23). BAF complexes modulate transcription through ATPase-dependent nucleosome sliding/ejection or the recruitment of other regulators (48, 64). They comprise three modules: a catalytic ATPase (BRG1/BRM), actin-related protein (ARP), and a base module that scaffolds the ATPase, nucleosome, and other regulators (Fig. 3A) (48). ARID1A or ARID1B are incorporated into the base module through a C-terminal Core Binding Region (CBR) made up of conserved CBRA and CBRB segments connected by intrinsically disordered loops (31).

**Figure 3:**
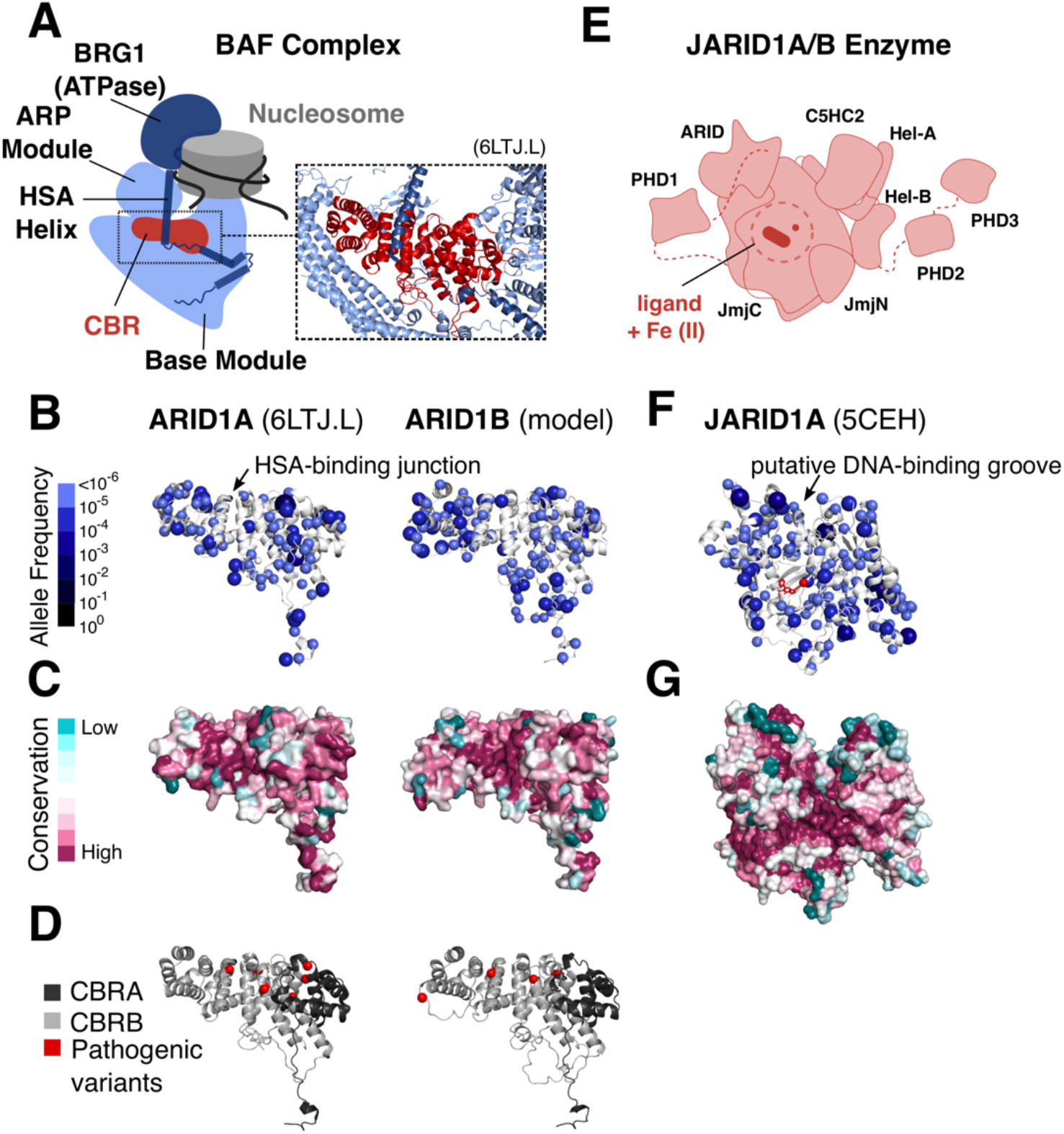
Validation of the 1D-to-3D approach. **(A)** Position of the ARID1A/1B CBR in the BAF complex. Solved structure of the ARID1A CBR (PDBid 6LTJ, chain L) and modelled structure of the ARID1B CBR annotated with missense variants **(B)**, sequence conservation in vertebrates **(C)**, and pathogenic variants **(D)**. **(E)** Schematic illustration of the JARID1A/1B enzymes (based on the PDB structures 5CEH and 5FUP). Solved structure of JARID1A annotated with missense variants (ligand shown in red) **(F)** and sequence conservation in metazoa **(G)**.

Mutations in the BAF complex promote tumorigenesis in multiple cancer types (reviewed in (20)). A recent analysis of missense cancer mutations in the BAF complex revealed that several mutations cluster at a junction between the ARID1A/B CBR and the helicase-SANT-associated (HSA) helix of the ATPase (Fig. 3A) (65). We find that the HSA-CBR junction is depleted of neutral missense variants (Fig. 3B), and this correlates well with evolutionary surface conservation of the CBR in vertebrates (Fig. 3C). We also found that pathogenic variants associated with Coffin-Siris Syndrome and non-syndromic intellectual disability variants map in proximity to the HSA-CBR junction (Fig. 3D). ARID1B is ranked as a top diagnostic gene for both Coffin-Siris syndrome and non-syndromic intellectual disabilities (66). ARID2 plays a functionally analogous role to the ARID1 family in the related Polybromo-associated BAF (PBAF) nucleosome remodeling complex (31). Even though ARID2 is distantly related to the ARID1 family, we found a similar depletion of missense mutations around the putative HSA helix binding site in a model of the ARID2 CBR domain (Fig. S1). The distribution of missense variants therefore serves as a valuable, additional layer of information for investigating key protein-protein interaction interfaces in multi-subunit complexes.

Next, we tested whether our approach could identify the catalytic site in members of the JARID1 subfamily. JARID1A/1B are Fe(II)- and 2-oxoglutarate (2-OG)-dependent dioxygenases that catalyse the demethylation of di- or tri-methylated histone H3K4 via their JmjC domain (Fig. 3E) (67). We report that the catalytic site in the JmjC domain is visibly missense depleted (Fig. 3F), correlating with evolutionary conservation (Fig. 3G). A similar depletion around the active site is observed for JARID1B (Fig. S2), but is less pronounced, consistent with the higher LOEUF and lower mZ metrics reported for JARID1B (Fig. 2A).

Apart from the catalytic site, depletion of missense variants in JARID1A is also observed in a groove between the ARID and C5HC2 domains, which is thought to accommodate doublestranded DNA (Fig. 3F; Movie S1) (49). The variants exhibit clear spatial surface segregation, with a lower abundance of variants on the face containing the catalytic site and higher abundance on the far surface of the enzyme (Movie S1). These findings validate our approach and demonstrate that missense variants complement the use of conservation data to identify surfaces involved in macromolecular interactions and enzyme active sites.

### Comparative Analysis of Domains Using the Vd/Vp Ratio

To further investigate the utility of missense variants at the domain level, we compared JARID1A and JARID1B. In addition to the JmjN-JmjC, ARID, and C5HC2 zinc finger domains, they also comprise three PHD fingers (Fig. 4A-B). Mapping missense variants on JARID1A and JARID1B sequences and calculating the Vd/Vp ratios for each domain revealed differences in missense variant depletion in each PHD finger (Fig. 4B). These differences may indicate paralog sub-functionalization at the level of individual domains. In JARID1A, PHD3 but not PHD1 is missense depleted, while in JARID1B, PHD1 but not PHD3 is missense depleted. Neither protein shows missense depletion in PHD2 (Fig. 4B).

**Figure 4:**
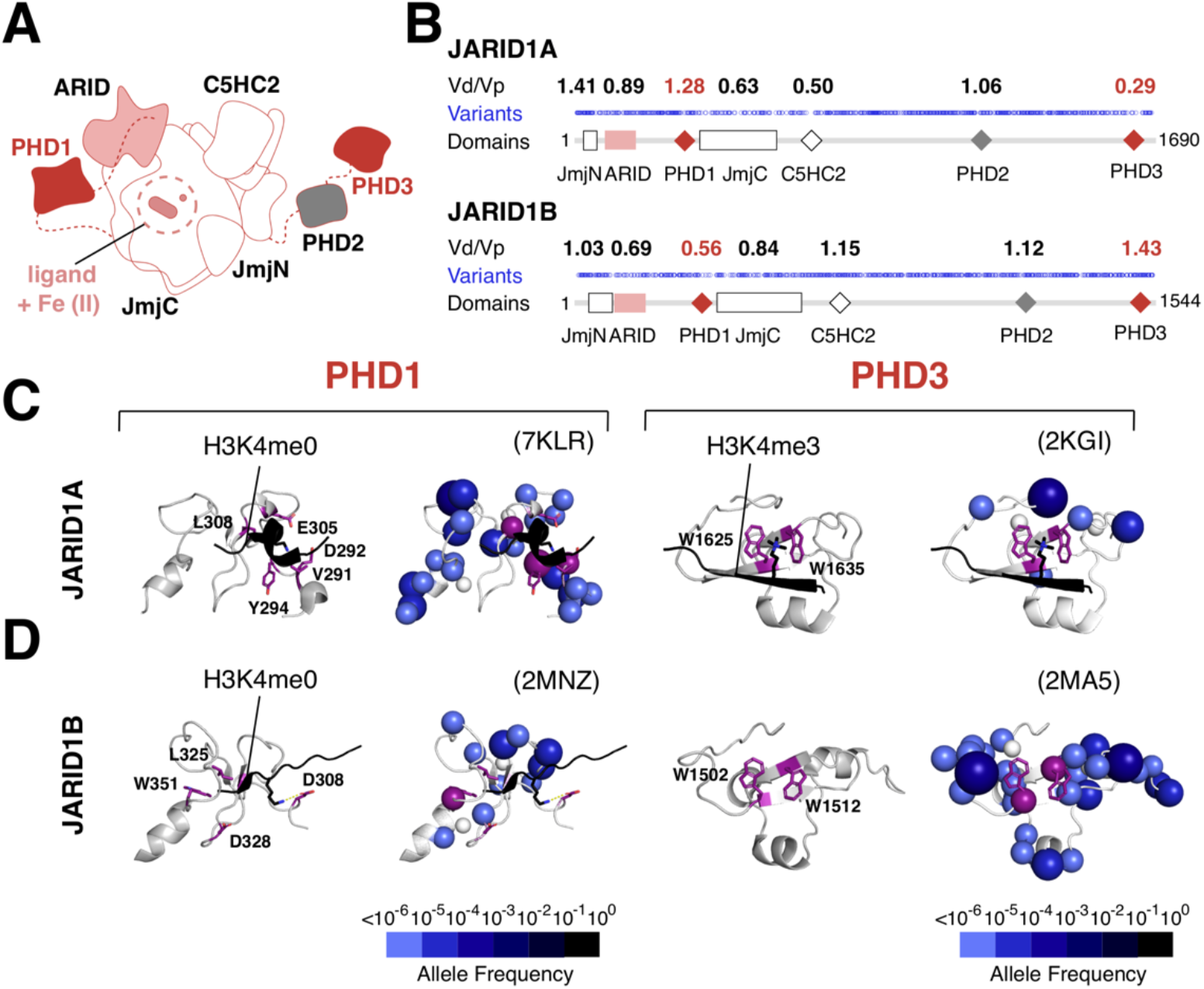
Comparative Analysis of Domains Using the Vd/Vp Ratio. **(A)** Schematic diagram of the JARID1A and JARID1B enzymes (based on PDB structures 5CEH and 5FUP). **(B)** 1D plots of missense variants in JARID1A/1B and with Vd/Vp ratios calculated for their functional domains, shown above the plot. **(C)** JARID1A and **(D)** JARID1B PHD1 and PHD3 domains shown without (left) and with (right) missense variants; histone peptides are shown in black, peptide-binding residues in purple, and zinc ions in white.

In JARID1A and JARID1B, PHD1 preferentially binds to unmethylated histone H3K4, leading to increased affinity of the JmjC domain for its methylated H3K4 substrates (68–70). In contrast, PHD3 is thought to be a reader of tri-methylated H3K4 marks (71, 72). These two domains tolerate sequence variation, suggesting that their ability to bind histone tails is not crucial for function. In particular, residues D292/Y294/L308 in JARID1A and W1502/W512 in JARID1B, which are required for histone peptide binding (72, 73), are affected by neutral missense variants (Fig. 4C-D). Conversely, residues responsible for histone peptide binding in PHD3 of JARID1A (Fig. 4C) and PHD1 of JARID1B (Fig. 4D) are under selective constraint, indicating that they are important in targeting or enhancing the activity of the JARID1 enzymes at their appropriate genomic locations. Consistent with their functional importance, PHD3 in JARID1A was shown to be critical in driving the oncogenic effects of a JARID1A-NUP98 fusion protein in acute myeloid leukemia (71). Furthermore, mutations in PHD1, but not PHD3 of JARID1B decreased the regulatory effects on cell migration in a model of triple negative breast cancer (72). In summary, functional differences in the PHD fingers correlate with different Vd/Vp ratio patterns in JARID1A and JARID1B, suggesting that the differences missense variant depletion we observe represent domain-level functional differences between the two paralogs.

### Limitations of the Vd/Vp Ratio

Despite its utility for comparing domains, it should be noted that Vd/Vp ratios are dependent on the user’s definition of domain boundaries. Where domain boundaries are not clearly defined, it is possible that Vd/Vp ratios might be over- or under-estimated. Furthermore, small sites that are missense depleted may also be overlooked. For example, the Vd/Vp ratios calculated for the chromobarrel domain of the ARID4 subfamily proteins are relatively high, yet mapping missense variants in 3D reveals depletion of variants in a conserved Tyr-Tyr-Trp-Tyr aromatic cage (Fig. 5).

**Figure 5:**
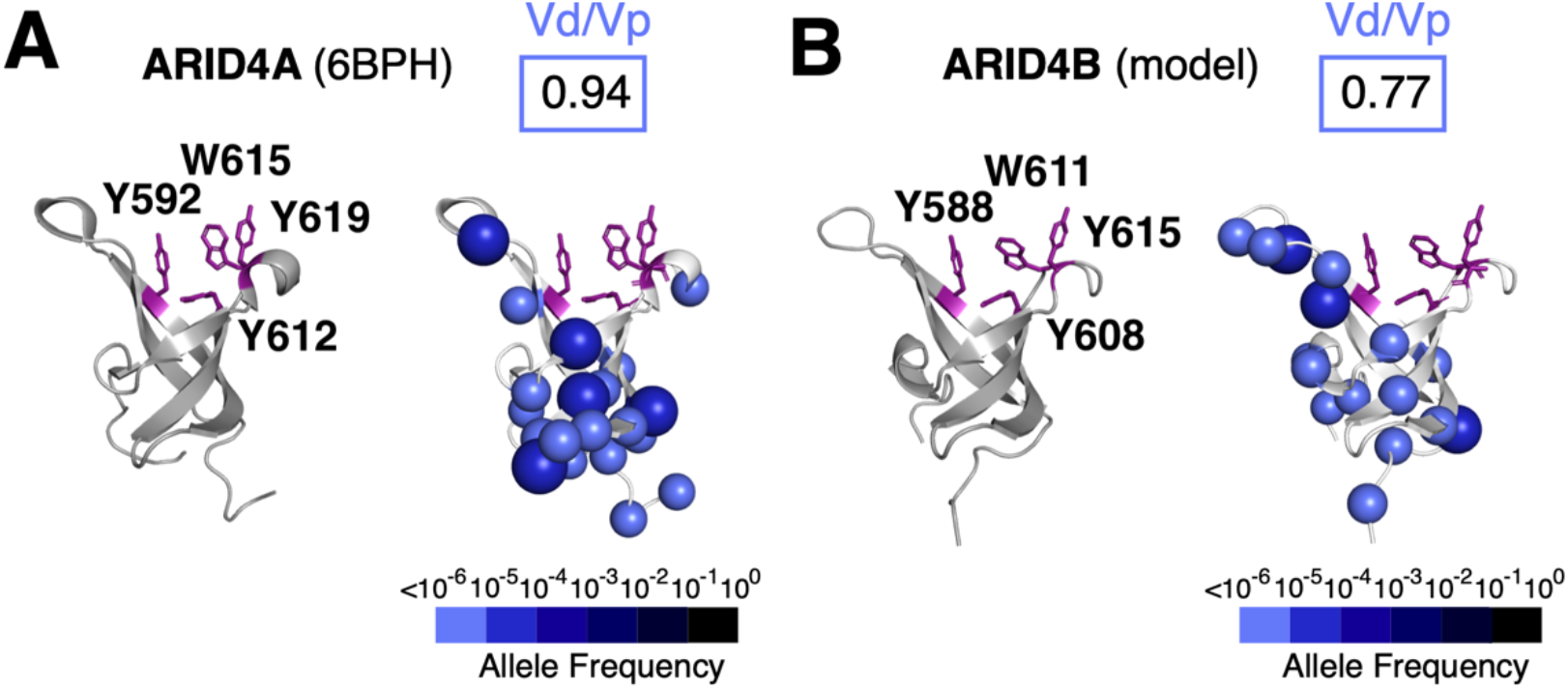
Limitations of the Vd/Vp Ratio. The ARID4A **(A)** and ARID4B **(B)** chromobarrel domains shown without (left) and with (right) missense variants. Putative histone methyl markbinding residues are indicated in purple.

The ARID4 subfamily comprises ARID4A and ARID4B (Fig. 2A), two paralogous, multidomain adapter proteins that recruit transcriptional regulators such as the retinoblastoma protein, androgen receptor, and the mSin3A histone deacetylase complex to gene promoters (38, 74). Given the presence of an aromatic cage, the chromobarrel domain was hypothesized to bind histone methyl marks (37). However, independent NMR and ITC titration experiments with the ARID4A chromobarrel domain and methylated histone peptides produced conflicting results (37, 75).

Mapping of missense variants shows that residues that form the aromatic cage are depleted of missense variants in both the solved structure of the ARID4A chromobarrel domain (Fig. 5A) and a model of the ARID4B chromobarrel domain (Fig. 5B). This suggests that methyl-lysine recognition is intact. This observation highlights the importance of mapping missense variants onto 3D structures, especially in the case of small domains, where the Vd/Vp ratio may not provide sufficient resolution to detect functionally important sites.

### Comparison of Missense Depletion Between Paralogs Indicates Sites of Sub-Functionalization

ARID4A and ARID4B have diverged only in vertebrate lineages (45) and share identical domain architecture (Fig. 2A). They participate in the same molecular pathways, including the recruitment of the mSin3A repressive complex to gene promoters (76) and co-activation of the androgen receptor in regulation of male fertility (74). However, ARID4A knockout mice are viable whereas ARID4B knockouts show early embryonic lethality (77). Moreover, ARID4B is necessary for spermatogenesis while ARID4A is not (74, 78). These findings indicate that of the two paralogs, ARID4B is likely a more critical determinant of cell fate decisions and cell cycle progression.

Since all domains of ARID4A and ARID4B are structurally related, we investigated if missense depletion could reveal functional differences between the two paralogs. We found domain-wide missense depletion in 1D to be more pronounced in ARID4B (Fig. S3) and observed a difference in the 3D distribution of missense variants on their hybrid Tudor domains (HTDs). The HTDs comprise two sub-domains HTD-1 and HTD-2 (Fig. 6A). Unlike structurally analogous Tudor domains, ARID4 HTDs lack an aromatic cage associated with histone binding in HTD-2 and instead are reported to exhibit DNA binding through HTD-1 (79, 80) (Fig. 6A). A conserved, structurally important glycine of ARID4B is associated with a developmental disorder variant (Fig. 6A).

**Figure 6:**
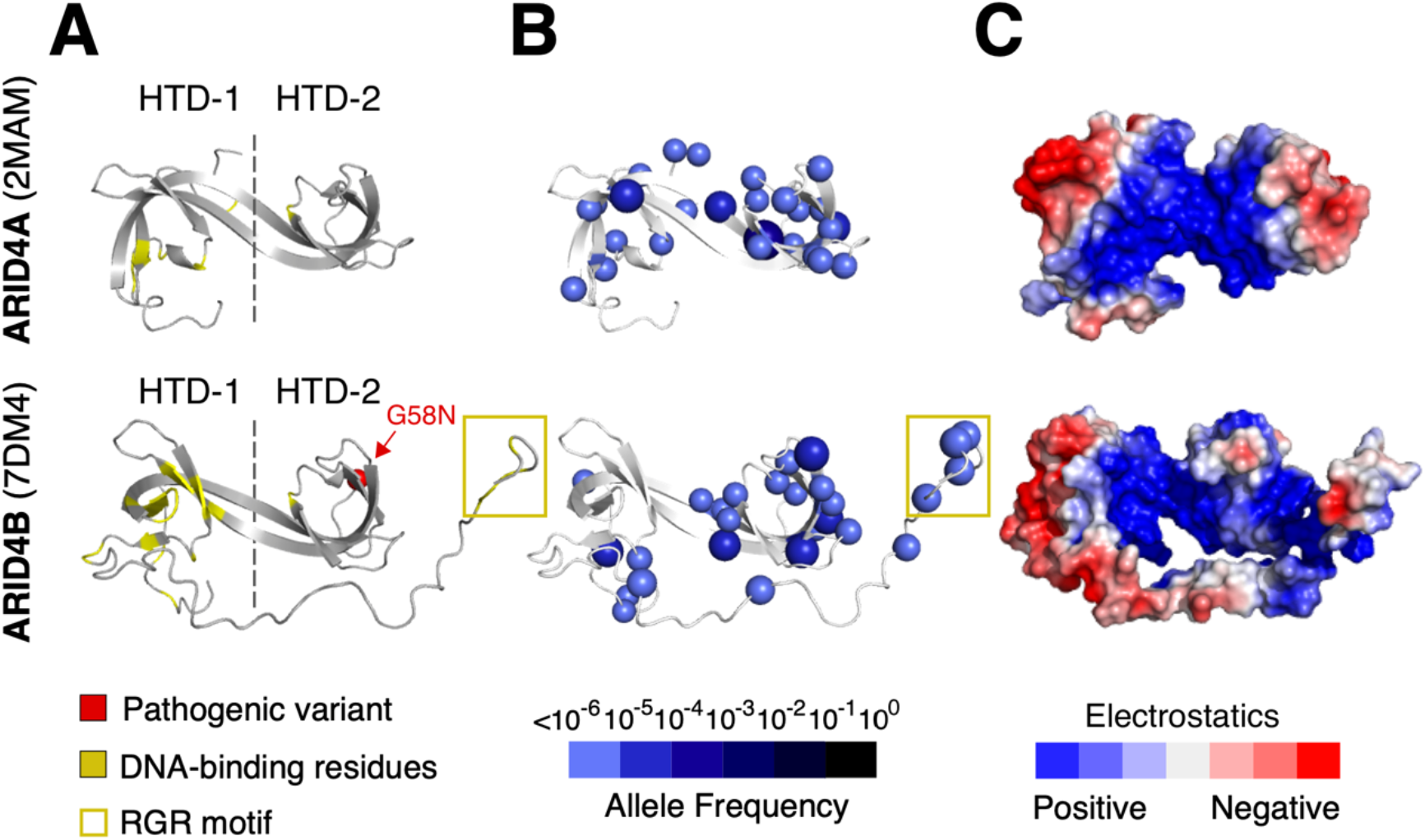
The ARID4 Subfamily. The ARID4A and ARID4B hybrid Tudor domain (HTD) shown with DNA binding residues **(A)**, missense variants **(B)**, and surface electrostatics **(C)**.

We find that HTD-1 of ARID4B is missense depleted, while HTD-1 of ARID4A is not (Fig. 6B). The depleted site corresponds to previously identified DNA-binding residues (Fig. 6B) and a positively-charged DNA-binding surface of the domain (Fig. 6C)(80). We also note that an RGR motif, recently found to enhance the DNA-binding affinity of ARID4B (80), tolerates missense variation (Fig. 6A). Overall, our findings indicate that the ARID4B Tudor domain likely contributes to the functional differences in DNA binding between ARID4A and ARID4B.

We next investigated if missense variants could also reveal paralog sub-functionalization in the ARID3 subfamily. The ARID3 proteins are transcription factors comprising the ARID domain and a C-terminal oligomerization domain called REKLES (Fig. 2A). We note that the Vd/Vp ratios of all three REKLES domains are > 1.00 (Fig. S4). While this suggests that the oligomerization is less likely to be required for ARID3 activity, these domains are short motifs with no available structural data, so may not be suitable for this analysis.

The ARID domain binds to AT-rich promoter sequences and is essential for ARID3 protein function (35, 81). In *Drosophila* and mice, knock out of the ARID3 orthologs, Dead ringer or Bright, respectively, is lethal (35, 82). In humans, the ARID3 subfamily has three members, where the ARID3A ARID domain shares 70 % sequence identity with the ARID domain of ARID3B and 87% identity with the ARID domain of ARID3C. Given this high degree of sequence identity, we hypothesized that differences in missense variation likely reflect paralog sub-functionalization.

The ARID domain is built from six core helices (H1-H6) and two larger loops L1 (between helices H1-H2) and L2 (between helices H4-H5) (Fig.7A). A solution structure of the *Drosophila* Dead ringer in complex with DNA (83) and NMR titration experiments of the ARID1A (84), ARID5B (85), and JARID1A (86) ARID domains suggest a common mode of DNA binding. This includes non-sequence specific contacts with the phosphate backbone by L1 and sequence-specific contacts with the major groove by a non-canonical helix-turn-helix motif formed by H4-L2-H5 (83–86)(Fig. 7B).

**Figure 7.**
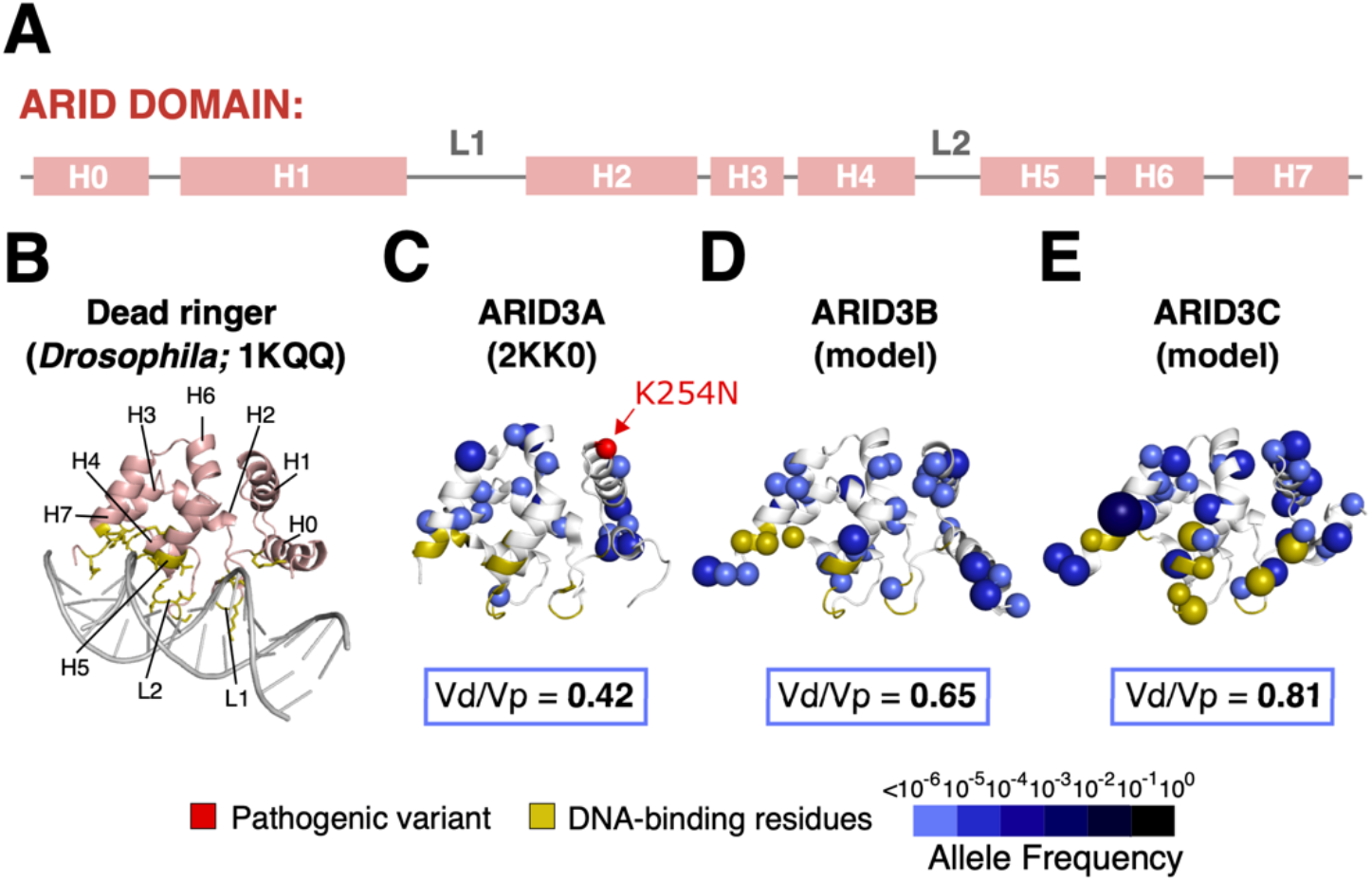
The ARID3 Subfamily. **(A)** Secondary structure of the ARID domain where helices are denoted with H and loops are denoted with L. The ARID3 subfamily has two additional flanking helices H0 and H7. **(B)** Solution structure of the Dead ringer ARID domain in complex with DNA. Human ARID3A **(C)**, ARID3B **(D)**, and ARID3C **(E)** ARID domains shown with missense variants.

The Vd/Vp ratios of ARID domains of the three human ARID3 paralogs are lowest in ARID3A and highest in ARID3C (Fig. 7C-E). In ARID3A, the proposed DNA binding residues in L1 and H4-L2-L5, inferred from sequence conservation with Dead ringer, are clear of missense variants. The domain also harbors a developmental disorder variant (59), supporting its functional importance (Fig. 7C). ARID3B has a higher Vd/Vp ratio and some missense variants map to residues typically involved in DNA binding in the ARID family. This suggests that DNA binding activity is less likely to be of functional importance in this paralog (Fig. 7D). Similarly, in ARID3C, several residues typically involved in DNA-binding are found to have reported variants (Fig. 7E). Missense variation can therefore also be leveraged to filter structurally similar domains in paralogs for functional importance.

### The ARID5B BAH Domain Shows Likely Loss and Gain of Function

Finally, we used 1D-to-3D to give insights on novel structure-guided functional predictions in ARID5B. ARID5B is a highly constrained gene (Fig. 2A) and a key regulator of liver metabolism, chondrogenesis, and adipogenesis (36, 87, 88). ARID5B is known to target the H3K9me2 demethylase PHF2 to gene promoters via its ARID domain (26, 36).

ARID5B has two isoforms, 1 and 2 (Fig. 8A). In *Xenopus*, isoform expression is spatially segregated during embryonic development, where isoform 1 shows higher abundance than isoform 2 (89). Isoform 1 also shows higher expression in healthy adult human tissues (57). Isoform1 has an additional N-terminal region that is highly conserved in a subset of vertebrate species (Fig. S5) and predicted to form a bromo-adjacent homology (BAH) domain (46, 90, 91); this likely defines the functional differences between the isoforms. We compared an AlphaFold model of the ARID5B BAH domain to experimentally determined BAH domain structures and used 1D-to-3D to map missense variants onto the domain.

**Figure 8:**
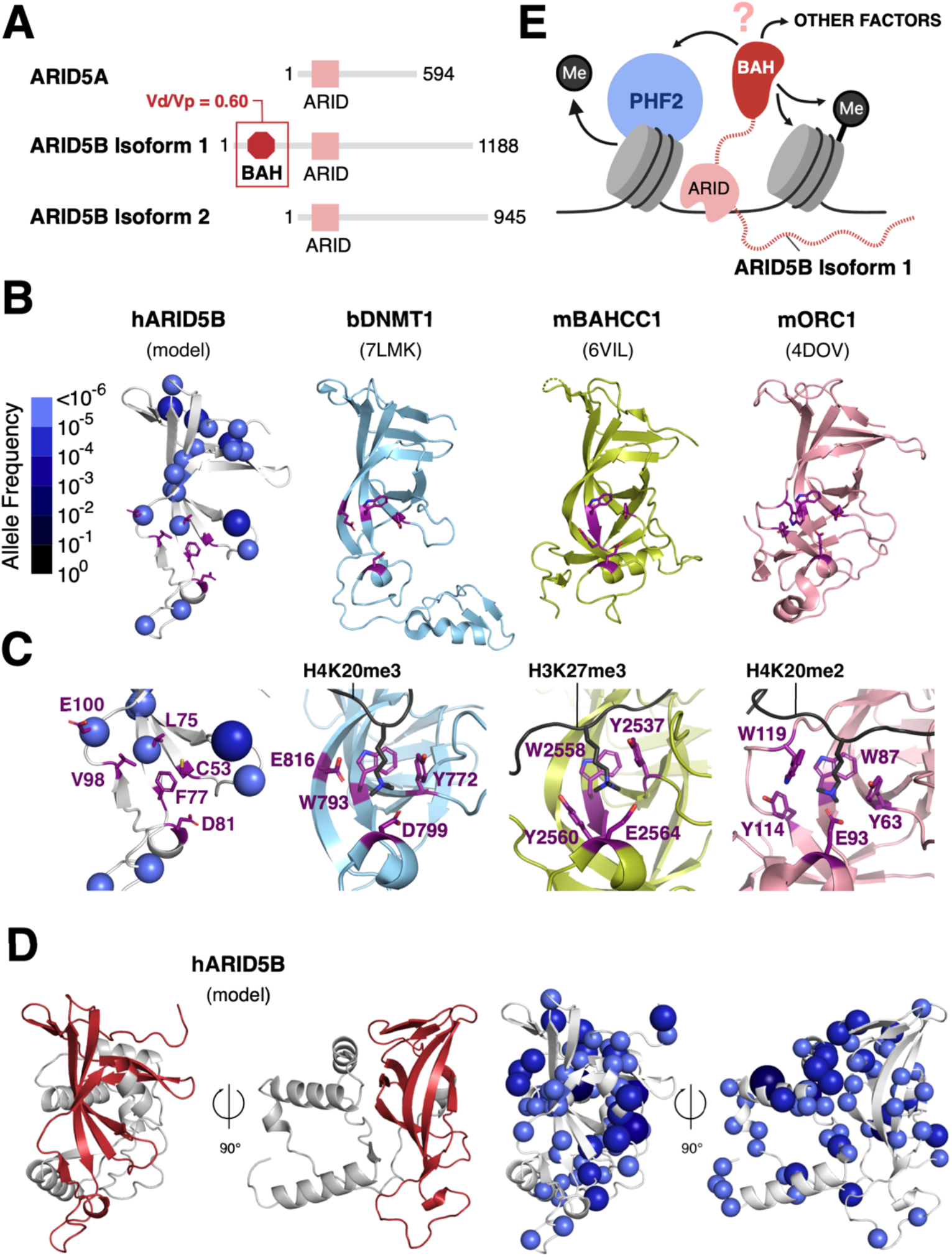
Loss and gain of function in ARID5B. **(A)** Domain architecture of the human ARID5 subfamily. **(B)** Variant-annotated model of the human ARID5B BAH domain compared to solved structures of the bovine DNMT1, mouse BAHCC1, and mouse ORC1 BAH domains (histone methyl mark-binding residues shown in purple). **(C)** A closer view of the methyl mark-binding sites (methylated histone peptides shown in black). **(D)** AlphaFold prediction of the ARID5B BAH domain with a C-terminal extension: two orientations, related by a 90 degree rotation, of the BAH domain (red) and the extension (grey) are shown in the left panel. Corresponding missense variants are shown on the same models in the right panel. **(E)** Potential binding interactions of the ARID5B BAH domain.

We identified bovine DNMT1, mouse BAHCC1, and mouse ORC1 BAH domains as the closest structural homologs to ARID5B (Fig. S6). All three homologs read histone methyl marks (Fig. 8B) (92–94). The lower lobe of each BAH fold contains a conserved aromatic cage and acidic residues that bind to methylated lysine through cation-pi and electrostatic/hydrogen bonding interactions, respectively (Fig. 8C). The lower lobe of the ARID5B BAH domain does not have an aromatic cage: one aromatic residue, F77, is present but two positions normally occupied by aromatic residues are replaced with small hydrophobic residues (C53 and L75). Two acidic residues, D81 and E100, are present but both E100 and the aliphatic L75 tolerate missense variation (Fig. 8C). Collectively, these amino acids changes, compared with classic BAH domains, and their tolerance of variation suggest that ARID5B BAH domain is unlikely to bind methyl-lysine marks at this site.

The AlphaFold prediction for the ARID5B BAH domain differs from those in DNMT1, BAHCC1 and ORC1 in that an additional, conserved segment C-terminal to the BAH domain forms part of the fold of this domain (Fig. 8D). When mapped to this extended AlphaFold model, we note that the missense variants are depleted on the highly conserved, positively-charged C-terminal helix, rather than the classic peptide binding interface of BAH domains. This suggests that the C-terminal domain extension could serve as a protein-protein or protein-nucleic acid interaction module within a chromatin context (Fig. 8E). These predictions call for experimental evidence. However, our analysis demonstrates the power of missense variation in screening for functional features together with structural data.

## Conclusion

We provide a convenient set of tools for mapping missense variants onto primary and tertiary structures of proteins. This approach allowed us to visually locate regions of proteins that are depleted of population variants, indicative of negative selection pressure. Using this approach, we demonstrated that mapping missense variants onto 3D structures in the context of a large family of proteins reveals functional insights. This method is complementary to phylogenetic conservation analysis and could be useful where insufficient phylogenetic data are available, for example when analyzing recent paralogs.

Using Vd/Vp ratios for individual domains may have a particular utility for researchers working on multidomain proteins where the goal is to identify which domains contribute essential functions. Ranking by Vd/Vp ratio could help prioritize which domains to delete in functional assays. This approach could also be useful for researchers seeking to provide minimal functional constructs of a protein for gene therapy approaches, where limiting the length of the protein, and therefore its coding sequence, can be critical for packaging into a virus (95–97).

Some limitations are noted. First, the sample size of the gnomAD dataset does not achieve mutational saturation (1). This sets limits on interpreting constraint in smaller domains/linear motifs and prevented us from analyzing proteins encoded on the sex chromosomes. Second, while our approach allows allele frequency to be visualized, it does not normalize for codon mutability, where nucleotide sequence composition skews missense enrichment in protein sequences. For example, methylated CpG dinucleotides are known to be hypermutable, resulting in an over-representation of variants in CpG rich and/or heavily methylated codons (98). Finally, some of our analyses were based on calculated models rather than experimentally-determined structures. As missense mapping depends on positional information, validation of these models is essential to confirm any interpretations (99).

Our findings build on previous studies showing that depletion of missense variants can serve to identify functionally important protein sites (7, 15). We demonstrate how 1D and 3D mapping approaches complement existing findings, provide context to understand the impact of pathogenic variants, functionally differentiate structurally similar domains in paralogs, and support formulation of novel mechanistic hypotheses. Although we focused on proteins with catalytic, DNA binding and epigenetic roles, our approach is applicable to a broad range of human protein functions.

## Acknowledgements

The authors would like to thank Sander Granneman and Philipp Voigt for critical reading of the manuscript. This work was supported by a Wellcome Trust with a Senior Research Fellowship to AGC (200898) and a Wellcome Centre core grant (203149). We thank Edward Wallace for help with phylogenetic analysis. We thank Hannah Wapenaar and Hayden Burdett for help with testing code.

## Author contributions

AGC conceived of and directed the research. GD executed research, developed metrics and wrote the code. GD and AGC co-wrote the MS.

## Data are available at

University of Edinburgh GitLab: https://git.ecdf.ed.ac.uk/cooklab/deak

University of Edinburgh DataShare: https://doi.org/10.7488/ds/3190.

## Supplementary figures

**Figure S1:**
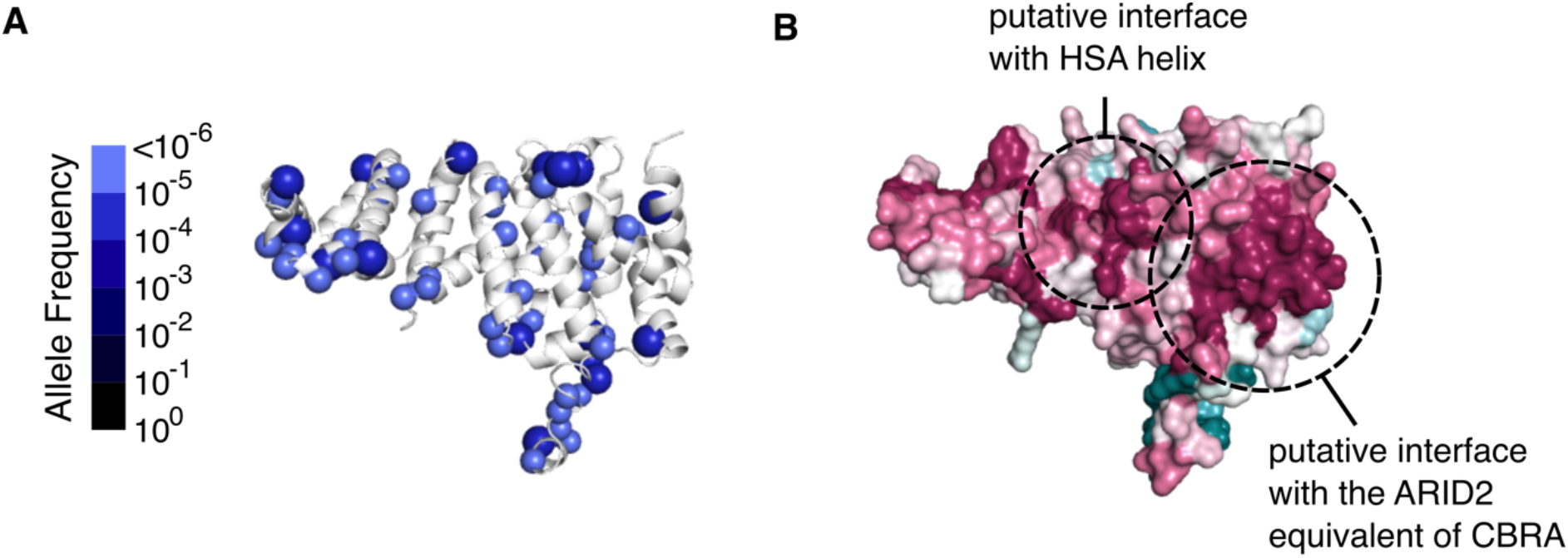
Modelled structure of the ARID2 equivalent of the ARID1A/1B CBRB (annotated with missense variants **(A)** and sequence conservation in metazoa **(B)**

**Figure S2:**
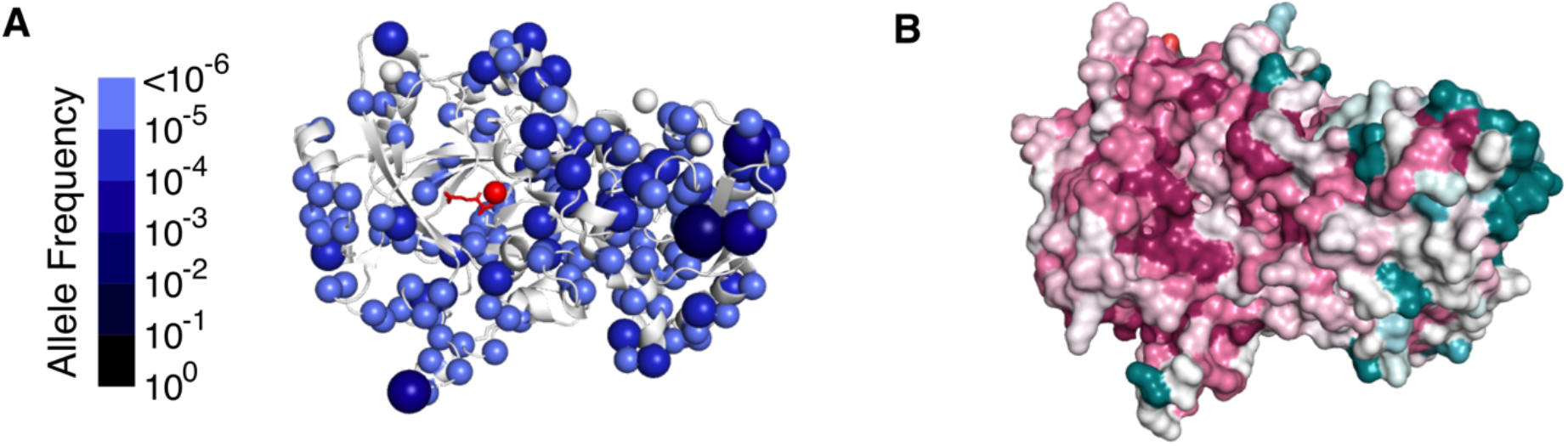
Solved structure of JARID1B (5FUP) annotated with missense variants **(A)** and sequence conservation in metazoa **(B)**

**Figure S3:**
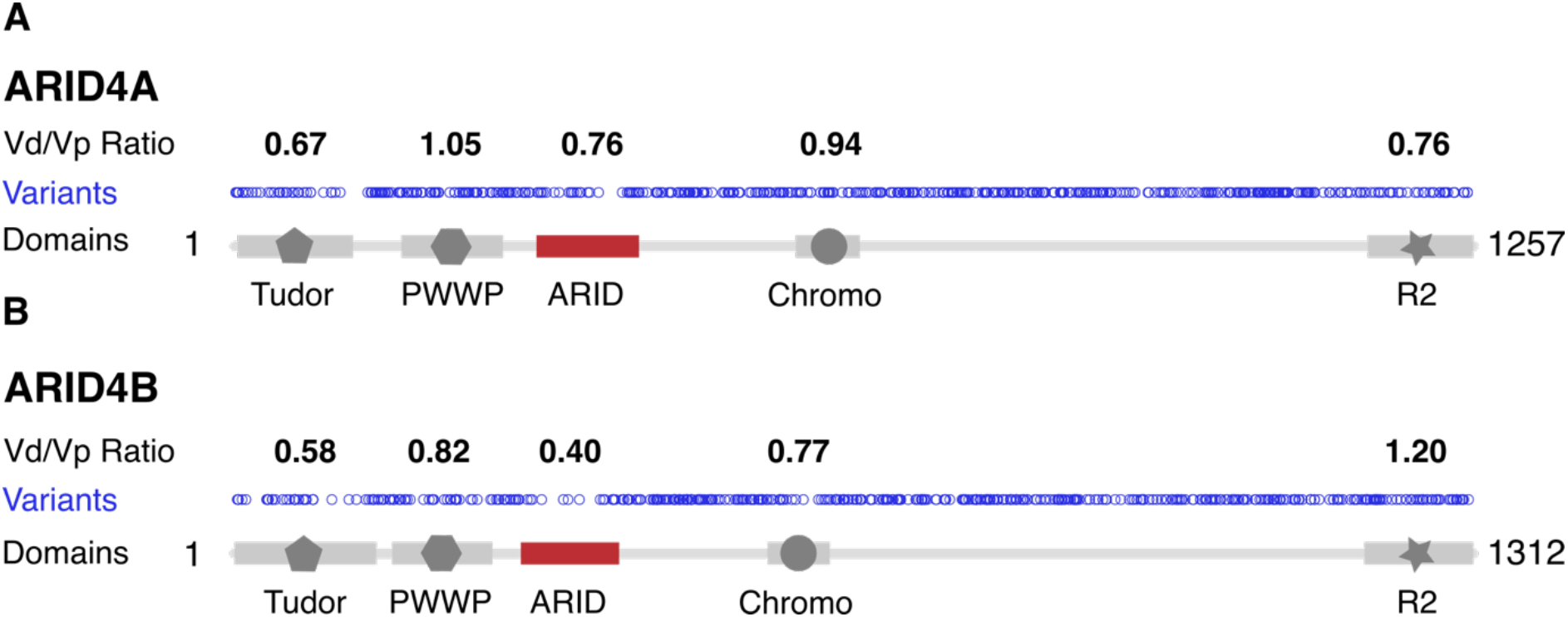
1D plots of missense variants in ARID4A **(A)** and ARID4B **(B)** and the Vd/Vp ratios calculated for their functional domains

**Figure S4:**
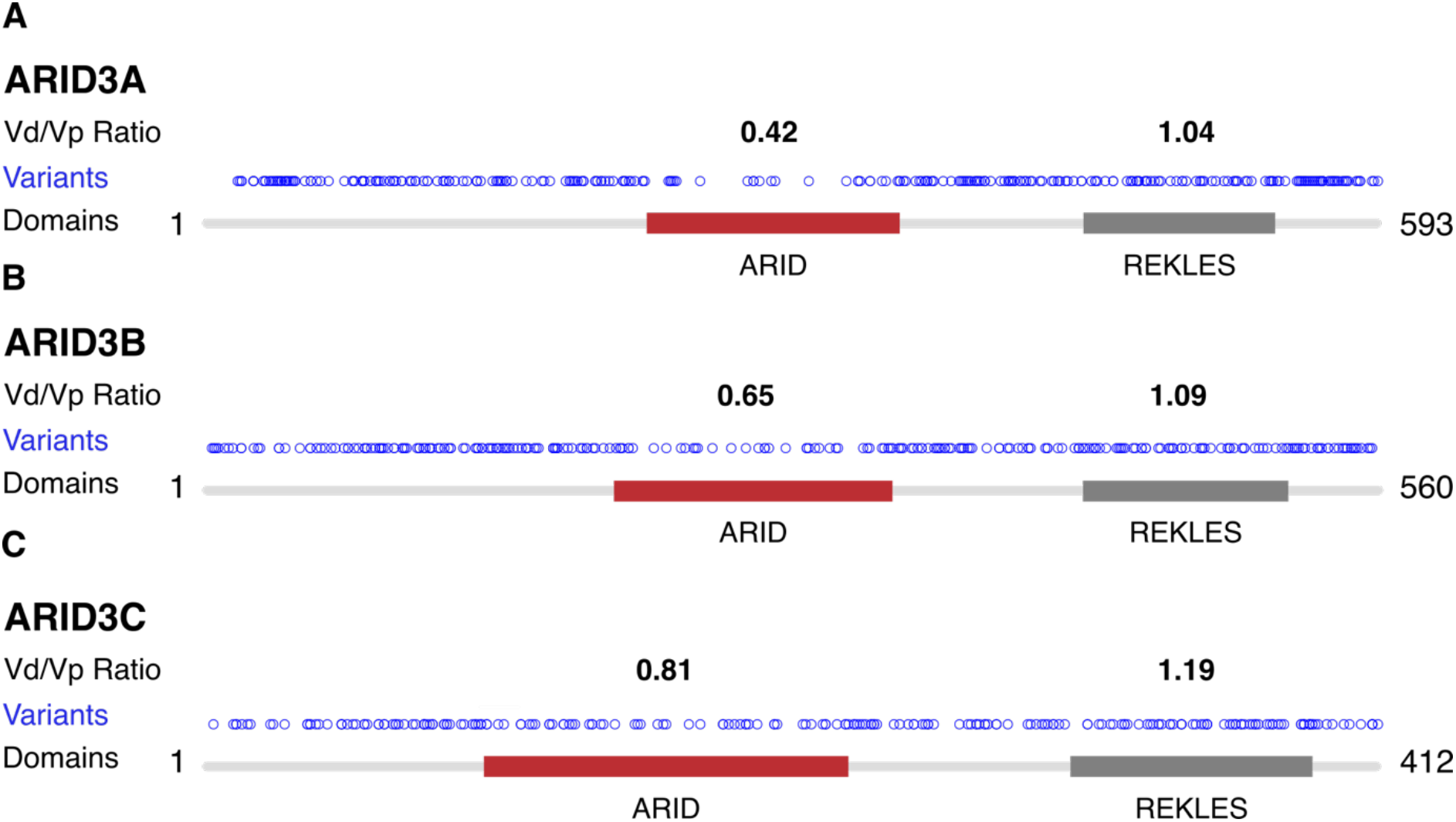
1D plots of missense variants in ARID3A **(A)**, ARID3B **(B)**, and ARID3C **(C)** and the V_d_/V_p_ ratios calculated for their functional domains

**Figure S5:**
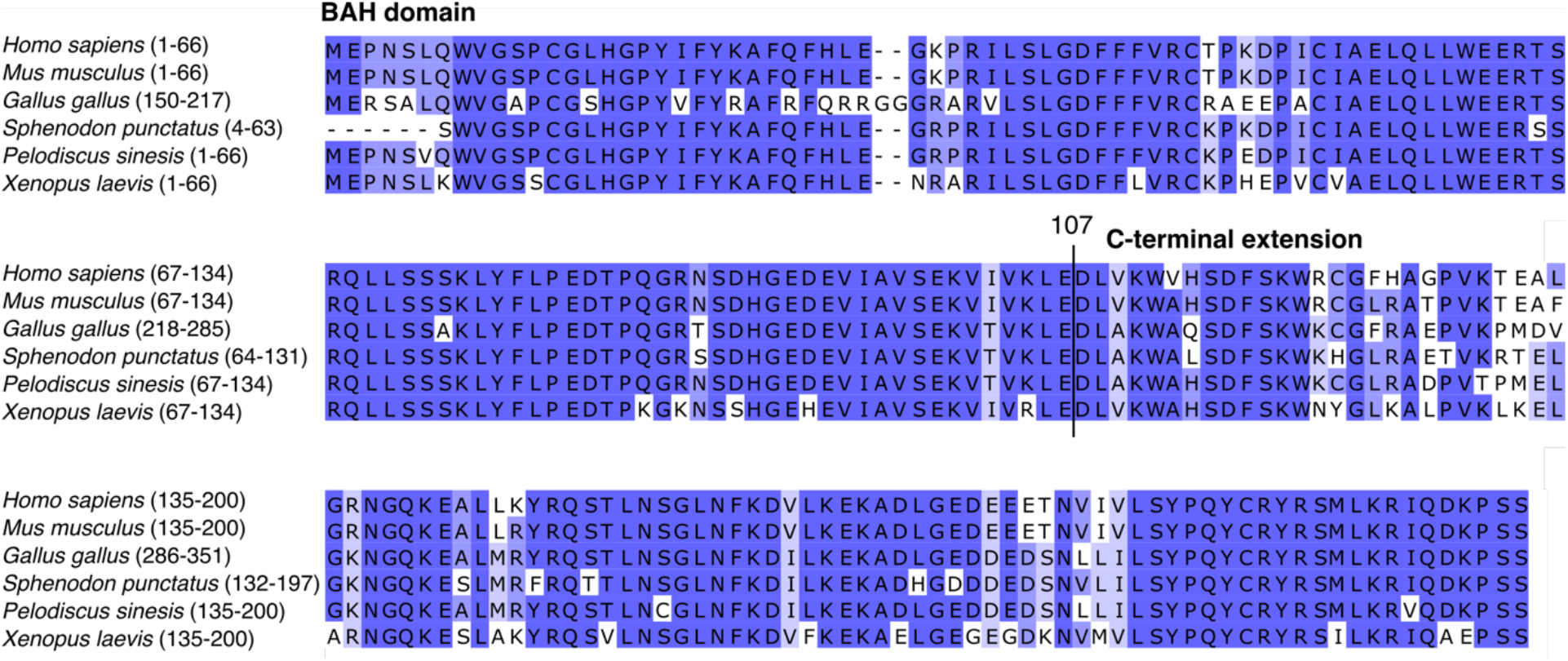
A vertebrate multiple sequence alignment of the N-terminal region of ARID5B isoform 1; The end of the BAH domain and the start of the C-terminal extension is marked by a vertical line at human aa 107. Darker blue represents higher percentage identity.

**Figure S6:**
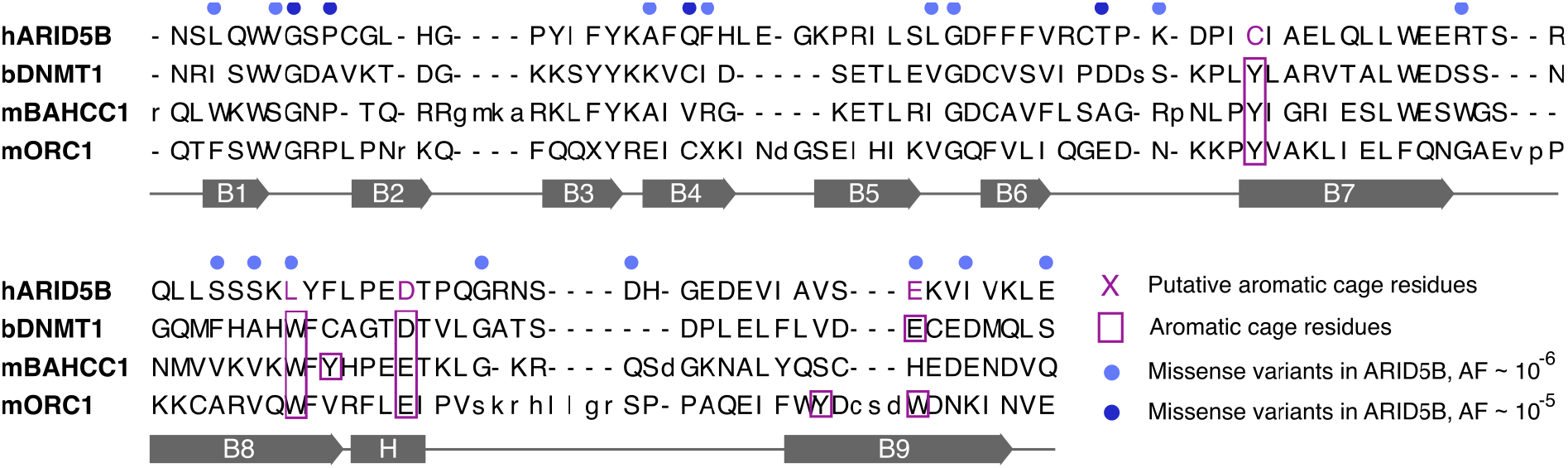
Structural alignment of the ARID5B BAH domain; h = human, b = bovine, m = murine

